# Learning Immune-Defectives Graph through Group Tests

**DOI:** 10.1101/015149

**Authors:** Abhinav Ganesan, Sidharth Jaggi, Venkatesh Saligrama

## Abstract

This paper deals with an abstraction of a unified problem of drug discovery and pathogen identification. Here, the “lead compounds” are abstracted as inhibitors, pathogenic proteins as defectives, and the mixture of “ineffective” chemical compounds and non-pathogenic proteins as normal items. A defective could be immune to the presence of an inhibitor in a test. So, a test containing a defective is positive iff it does not contain its “associated” inhibitor. The goal of this paper is to identify the defectives, inhibitors, and their “associations” with high probability, or in other words, learn the Immune Defectives Graph (IDG). We propose a probabilistic non-adaptive pooling design, a probabilistic two-stage adaptive pooling design and decoding algorithms for learning the IDG. For the two-stage adaptive-pooling design, we show that the sample complexity of the number of tests required to guarantee recovery of the inhibitors, defectives and their associations with high probability, i.e., the upper bound, exceeds the proposed lower bound by a logarithmic multiplicative factor in the number of items. For the non-adaptive pooling design, in the large inhibitor regime, we show that the upper bound exceeds the proposed lower bound by a logarithmic multiplicative factor in the number of inhibitors.

## I. Introduction

Drug discovery is known to be a tedious process, where early stages of drug discovery involve finding ‘blocker’ or ‘lead’ compounds amidst billions of chemical compounds [1], [2]. These lead compounds bind to a biomolecular target, which is a pathogenic protein, and thus inhibit the function of the protein. Such compounds are later used to produce new drugs. Finding a relatively small number of lead compounds amidst a large collection of chemical compounds is a challenging task. A complementary problem involves identifying pathogenic proteins amidst non-pathogenic ones, both of which are structurally identical in some respects. For instance, out of five known species of ebolavirus, only four of them are pathogenic to humans (see p. 5 in [2]) and a similar example can be found in arenavirus [3]. Some of these pathogenic proteins might share a common inhibitory mechanism against a lead compound which serves to distinguish them from the non-pathogenic ones [3]. So, finding potential pathogenic proteins amidst a large collection of biomolecules by testing them against known inhibitory compounds is a problem complementary to the problem of lead compound discovery. The lead compounds can be abstracted as inhibitor items, the pathogenic proteins as defective items, and the others as normal items. Now, the above problems can be combined to be viewed as an inhibitor-defective classification problem on the mixture of pathogenic and non-pathogenic proteins, and billions of chemical compounds. This unifies the process of finding both the pathogenic proteins and the lead compounds. An efficient means of solving this problem could potentially be applied in computer-assisted drug and pathogen identification. A natural consideration is that, while some pathogenic proteins might be inhibited by some lead compounds, other pathogenic proteins might be immune to some of these lead compounds present in the mixture of items. In other words, each defective item is possibly immune to the presence of some inhibitor items so that its expression cannot be prevented by the presence of those inhibitors when tested together. By definition, an inhibitor inhibits at least one defective. Learning this inhibitor-defective interaction as well as classifying the inhibitors and defectives efficiently through group testing is presented this work.

A representation of this model, which we refer to as the Immune-Defectives Graph (IDG) model, is given in Fig. 1. The presence of a directed edge between a pair of vertices 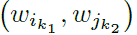 represents the inhibition of the defective 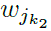 by the inhibitor 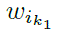 and the absence of a directed edge between a pair of vertices 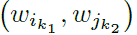 indicates that the inhibitor 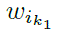 does not affect the expression of the defective 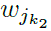 when tested together.

**Fig. 1.**
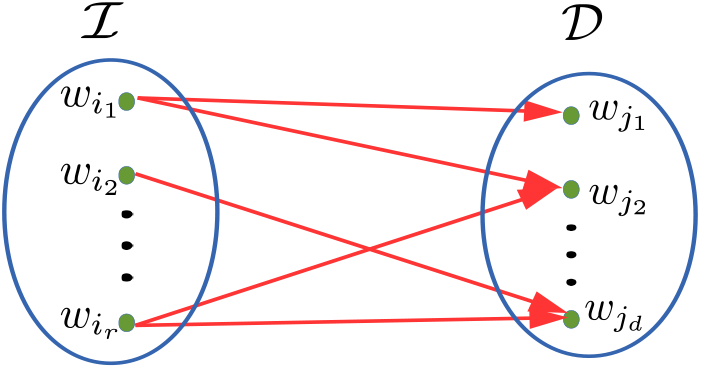
A representation of the IDG Model, where ℐ represents the set of inhibitors and 𝒟 represents the set of defectives.

***Example 1:*** An instance of the IDG model is given in Fig. 2. In this example, the outcome of a test is positive iff a defective 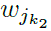, for some *k*_2_, is present in the test and its associated inhibitor 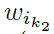 does not appear in the test. Observe that if the item-pair 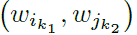, for *k*_1_ ≠ *k*_2_, appears in a test and 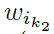 does not appear in the test, then the outcome is positive. Also, if the item-pair 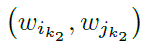 appears in a test and if 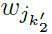 also appears in the test but not 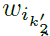, then the test outcome is positive. But if the appearance of every defective 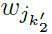 in a test is compensated by the appearance of its associated inhibitor 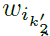 in the test, then the test outcome is negative. The outcome of a test is also negative when none of the defectives appear in a test.

**Fig. 2.**
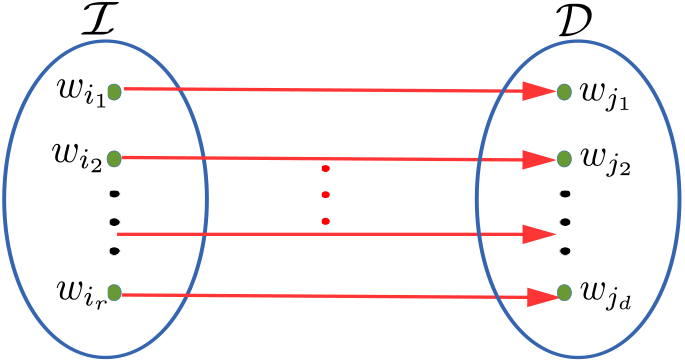
An example for the IDG Model where each defective is associated with a single distinct inhibitor so that *r* = *d*.

The IDG model can also be viewed as a generalization of the 1-inhibitor model introduced by Farach et al. in [4]. This model was motivated by errors in blood testing where blocker compounds (i.e., inhibitors) block the expression of defectives in a test [5]. This is also motivated by drug discovery applications where the inhibitors are actually desirable items that inhibit the pathogens [6]. In the 1-inhibitor model, a test outcome is positive iff there is at least one defective and no inhibitors in the test. So, the presence of a single inhibitor is sufficient to ensure that the test outcome is negative.

Efficient testing involves pooling different items together in a every test, in other words, called group testing [7] so that the number of tests can be minimized. The pooling methodology can be of two kinds, namely non-adaptive and adaptive pooling designs. In non-adaptive pooling designs, any pool constructed for testing is independent of the previous test outcomes, while in adaptive pooling designs, some constructed pools might depend on the previous test outcomes. A *k*-stage adaptive pooling design is comprised of pool construction and testing in *k*-stages, where the pools constructed for (non-adaptive) testing in the *k*^th^ stage depend on the outcomes in the previous stages. While adaptive group testing generally requires lesser number of tests than non-adaptive group testing, the latter inherently supports parallel testing of multiple pools. Thus, non-adaptive group testing is more economical (because it allows for automation) as well as saves time (because the pools can be prepared all at once) which are of concern in library screening applications [8]. The 1-inhibitor model has been extensively studied, and several adaptive and nonadaptive pooling designs for classification of the inhibitors and the defectives are known (refer, [9]–[12]). A detailed survey of known non-adaptive and adaptive pooling designs for the 1-inhibitor model is given in [13]. The best (in terms of number of tests) known non-adaptive pooling design that guarantees high probability classification of the inhibitors and defectives is proposed in [13]. The non-adaptive pooling design proposed in [13] requires *O*(*d* log *n*) tests in the *r* = *O*(*d*) regime and 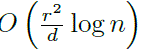 tests in the *d* = *o*(*r*) regime to guarantee classification of both the inhibitors and defectives with high probability^1^. In the small inhibitor, i.e., *r* = *O*(*d*) regime, the upper bound on the number of tests matches with the lower bound while in the large inhibitor, i.e., *d* = *o*(*r*) regime, the upper bound exceeds the lower bound of 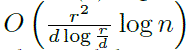 by a log 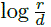 multiplicative factor. Nonetheless, the 1-inhibitor model constrains that every inhibitor must inhibit every defective, which is likely to be a tight requirement in practice. So, the IDG model is a more practical variant of the 1-inhibitor model.

A formal presentation of the IDG model and the goals of this paper are given in the next section.

***Notations:*** The Bernoulli distribution with parameter *p* is denoted by **ℬ**(*p*), where *p* denotes the probability of the Bernoulli random variable taking a value of one. The set of binary numbers is denoted by 𝔹. Matrices are indicated by boldface uppercase letters and vectors by boldface lowercase letters. The row-*i*, column-*j* entry of a matrix **M** is denoted by **M**(*i, j*), and the coordinate-*i* of a vector **y** is denoted by **y**(*i*). All the logarithms in this paper are taken to the base two.

## II. The IDG Model

Consider a set of items 𝒲 indexed as *w_i_*, …, *w_n_* comprised of *r* inhibitors, *d* defectives, and *n* – *r* – *d* normal items. It is assumed throughout the paper that *r*, *d* = *o*(*n*).

*Definition 1:* An item pair (*w_i_, w_j_*), for *i ≠ j*, is said to be *associated* when the inhibitor *w_i_* inhibits the expression of the defective *w_j_*. An item pair (*w_i_,w_j_*), for *i ≠ j*, is said to be *non-associated* if either the inhibitor *w_i_* does not inhibit the expression of the defective *w_j_* or if *w_i_* is not an inhibitor or if *w_j_* is not a defective.

In general, the mention of an item pair (*w_i_, w_j_*) need not mean that *w_i_* is an inhibitor and *w_j_* is a defective. This is understood from the context.

*Definition 2:* An *association graph* is a left to right directed bipartite graph **ℬ** = (ℐ, 𝒟, ℰ), where the set of vertices (on the left hand side) ℐ = {*w_i_1__, w_i_2__*, …, *w_i_r__*} ⊂ 𝒲 denotes the set of inhibitors, the set of vertices (on the right hand side) 𝒟 = {*w_j_1__, w_j_2__*, …, *w_j_d__*} ⊂ 𝒲 denotes the set of defectives, and ℰ is a collection of directed edges from ℐ to 𝒟. A directed edge *e* = (*w_i_,w_k_*) ∈ **ℰ**, for *i* ∈ {*i_1_, …, i_r_*}, *j* ∈ {*j_i_, …, j_d_*}, denotes that the inhibitor *w_i_* inhibits the expression of the defective *w_k_*.

We refer to ℰ (ℐ, 𝒟) *conditioned on the sets* (ℐ, 𝒟) to be the *association pattern* on (ℐ, 𝒟).

A pooling design is denoted by a test matrix **M** ∈ 𝔹^*T×n*^, where the *j*^th^ item appears in the *i*^th^ test iff **M** (*i, j*) = 1. A test outcome is positive iff the test contains at least one defective without any of its associated inhibitors. A positive outcome is denoted by one and a negative outcome by zero.

It is assumed throughout the paper that the defectives are not mutually obscuring, i.e., a defective does not function as an inhibitor for some other defective. In other words, the set of inhibitors ℐ and the set of defectives 𝒟 are disjoint.

The goal of this paper is to identify the association graph, or in informal terms, learn the IDG. Thus, the objectives are two-fold as represented by Fig. 3.

**Fig. 3.**
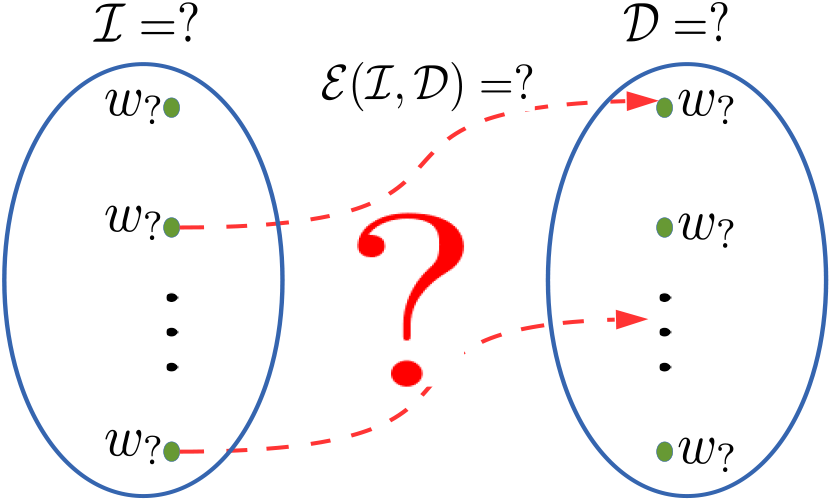
Here, the presence of a directed arrow represents an association between an inhibitor and a defective. The problem statement is to identify the set of inhibitors ℐ, defectives 𝒟 and the association pattern ℰ(ℐ,𝒟).

1) Identify all the defectives.
2) Identify all the inhibitors and also their association pattern with the defectives.

This problem is further mathematically formulated as follows. Denote the actual set of inhibitors, normal items, and defectives by ℐ, 𝒩, and 𝒟 respectively so that ℐ ∪︀ 𝒩 ∪︀ 𝒟 = 𝒲. The actual association pattern between the actual inhibitor and defective sets is represented by ℰ (ℐ, 𝒟). Let 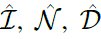, and 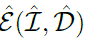 denote the declared set of inhibitors, normal items, defectives, and declared association pattern between 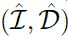 respectively. The target is to meet the following error metric.

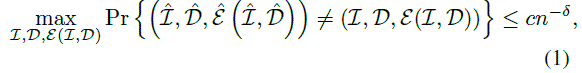

for some *c* > 0 and fixed ***δ*** > 0. We propose pooling designs and decoding algorithms, and lower bounds on the number of tests required to satisfy the above error metric. It is assumed that the defective and the inhibitor sets are distributed uniformly across the items, i.e., the probability that any given subset of *r + d* items constitutes all the defectives and inhibitors is given by 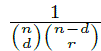 It is also assumed that the association pattern ℰ (ℐ, 𝒟) is uniformly distributed over all possible association patterns on (ℐ, 𝒟).

We consider two variants of the IDG model. The first being the case where the maximum number of inhibitors that can inhibit any defective, given by *I_max_*, is known. We refer to this model as the IDG with side information (IDG-WSI) model. For example, Fig. 2 represents a case where *I_max_* = 1. While it is known that *I_max_* = 1, it is unknown which among the items *w_i_*, …, *w_n_* represent which inhibitors and defectives. For a given value of (*r, d*), not all positive integer values of *I_max_* ≤ *r* might be feasible. For instance, if (*r, d*) = (3, 2), then *I_max_* = 1 is not feasible because, by definition, each inhibitor is associated with at least one defective. So, in the IDG-WSI model, we assume that the given value of *I_max_* is feasible for the (*r, d*) tuple. In particular, if (*c – 1)d < r ≤ cd* for some integer *c* ≥ 1, then *I_max_* ≥ *c*. This immediately follows from the fact that each inhibitor must be associated with at least one defective.

The other variant of the IDG model we consider in this paper is the case where there is no side information about the inhibitor-defective associations, which means that each defective can be inhibited by as many as *r* inhibitors. We refer to this model as the IDG-No Side Information (IDG-NSI) model. For both the models, the goals (as stated in the beginning of this section) are the same.

The contributions of this paper for the IDG models are summarized below.

- The sample complexity of the number of tests sufficient to recover the association graph while satisfying the error metric (1) using the proposed – non-adaptive pooling design is given by *T_NA_* = *O* ((*r* + *d*)^2^ log *n*) and *T_NA_* = *O* ((*I*_max_ + *d*)^2^ log *n*) tests for the IDG-NSI and IDG-WSI models respectively (Theorem 1, Section III). – two-stage adaptive pooling design is given by *T_NA_* = *O* (*rd* log *n*) and *T_A_ = O* (*I*_max_*d* log *n*) tests for the IDG-NSI and IDG-WSI models respectively (Theorem 2, Section III).
- In Section IV (Theorem 4 and Theorem 5), lower bounds of

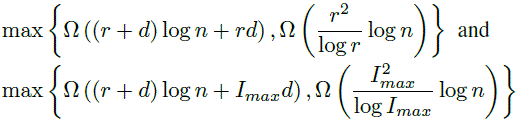

are obtained for non-adaptive pooling designs for the IDG-NSI and IDG-WSI models respectively. The first lower bound in both the models is valid for adaptive pooling designs also. Thus, the upper bounds on the number of tests for the proposed two-stage adaptive pooling design exceed the lower bounds by log *n* multiplicative factors in both the models. For the proposed non-adaptive pooling design, the upper bounds exceed the lower bounds by log *r* and log *I_max_* multiplicative factors in the *d = O(I_max_*) and *d = O(r)* regimes in the IDG-NSI and IDG-WSI models respectively.

*Extension of the results on the upper and lower bounds on the number of tests to the case where only upper bounds on the number of inhibitors (given by R) and defectives (given by D) are known instead of their exact numbers is straightforward. The target error metric in (1) is re-formulated as maximum error probability criterion over all combinations of number of inhibitors and defectives. The results for this case follow by replacing r by R and d by D in the upper and lower bounds on the number of tests.*

There are various generalizations of the 1-inhibitor model considered in the literature. These models are summarized in the following sub-section to show that the model considered in this paper, to the best of our knowledge, has not been studied in the literature.

### A. Prior Works

The 1-inhibitor model can be generalized in various directions, mostly influenced by generalizations of the classical group testing model. The various generalizations are listed below and briefly described. Though none of these generalizations include the model studied in this paper, it is worthwhile to understand the differences between these models and the IDG model.

A generalization of the 1-inhibitor model, namely *k*-inhibitor model was introduced in [14]. In the *k*-inhibitor model, an outcome is positive iff a test contains at least one defective and no more than *k* – 1 inhibitors. So, the number of inhibitors must be no less than a certain threshold *k* to cancel the effect of any defective. This model is different from the model introduced in this paper because, in the IDG model, a single associated inhibitor is enough to cancel the effect of a defective. Further, none of the inhibitors might be able to cancel the effect of a defective because the defective might not be associated with any inhibitor. A model loosely related with the 1-inhibitor model, namely mutually obscuring defectives model was introduced in [15]. Here, it was assumed that multiple defectives could cancel the effect of each other, and hence the outcome of a test containing multiple defectives could be negative. Thus, a defective can also function as a inhibitor. However, in this paper, the sets of defectives and inhibitors are assumed to be disjoint. The threshold (classical) group testing model is where a test outcome is positive if the test contains at least *u* defectives, negative if it contains no more than *l* defectives and arbitrarily positive or negative otherwise [16]. This model was combined with the *k*-inhibitor model and non-adaptive pooling designs for the resulting model was proposed in [17].

A non-adaptive pooling design for the general inhibitor model was proposed in [18]. Here, the goal was to identify all the defectives with no prior assumption on the cancellation effect of the inhibitors on the defectives, i.e, the underlying unknown inhibitor model could be a 1-inhibitor, *k*-inhibitor model, or even the ID model introduced in this paper. However, the difference from our work is that, we aim to identify the association graph or, in other words, the cancellation effect of the inhibitors also apart from identification of the defectives. But this cancellation effect does not include the *k*-inhibitor model cancellation effect as noted earlier. Group testing on complex model was introduced in [19]. In the complex model, a test outcome is positive iff the test contains at least one of the defective sets. So, here the notion of defectives items is generalized to sets of defective items called defective sets. This complex model was combined with the general inhibitor model and non-adaptive pooling designs for identification of defectives was proposed in [20]. Our work is different from [20] for the same reasons as stated for [18]. Group testing on bipartite graphs was proposed in [21] as a special case of the complex model. Here, the left hand side of the bipartite graph represents the bait proteins and the right hand side represents the prey proteins. It is known a priori which items are baits and which ones are preys. The edges in the bipartite graph represent associations between the baits and preys. A test outcome is positive iff the test contains associated items and the goal was to identify these associations. Clearly, this model is different from the IDG because, in the IDG model, there are three types of items involved and the interactions between the three types of items are different from that in [21].

In the next section, we propose a probabilistic non-adaptive and a probabilistic two-stage adaptive pooling design and decoding algorithms for both the variants of the IDG model.

## III. Pooling Designs and Decoding Algorithmg

In this section, we propose a non-adaptive pooling design and decoding algorithm as well as a two-stage adaptive pooling design and decoding algorithm for the IDG-WSI Model. The pooling designs and decoding algorithms for the IDG-NsI model follows from those for the the IDG-WSI Model by replacing *I_max_* by *r*.

*Non-adaptive pooling design*: The pools are generated from the matrix **M**_*NA*_ ∈ 𝔹*^T_NA×n_^*. The entries of **M***_NA_* are i.i.d. as ***ℬ** (p_1_).* Test the pools denoted by the rows of **M***_NA_*. Let the outcome vector be given by **y** ∈ 𝔹*^T_NA×1_^*. The exact value of *T_NA_* is specified in (11) and (12) (where, *T_NA_ = β_NA_* log *n*), and its scaling is given in Theorem 1. The exact value of *p_1_* is also given in Theorem 1.

*Adaptive pooling design:* A set of pools are generated from the matrix **M**_1_ ∈ 𝔹*^T_1_×n^* whose entries are i.i.d. as *ℬ*(*p*_1_). The pools denoted by the rows of **M**_1_ are tested first and all the defectives are classified from the outcome vector **y**_1_ ∈ 𝔹*^T_l_×1^*. Denote the number of items declared defectives by *d̂* and the set of declared defectives by {*û_1_, û_2_, …, û_d̂_*}, If *d̂ ≠ d* an error is declared. We keep these declared defectives aside and generate another pooling matrix **M**_2_ ∈ 𝔹*^T_2×(n‒d)_^*, whose entries are i.i.d. as ℬ(*p_2_*), for the rest of the items. Now, test the pools denoted by the rows of the matrix M_2_ along with each of the items declared defectives and the outcomes are denoted by **y***_̂u_1__*, **y***_̂u_2__* …, **y***_̂u_d__* ∈ 𝔹*^T_2×1_^*. The two stages of testing are done non-adaptively, and hence the pooling scheme is a two-stage adaptive pooling design. The exact values of *p*_1_ and *p*_2_ are given in Theorem 2. The scaling of *T*_1_ and *T*_2_ are also given in Theorem 2 and their exact values are given in (11) and (13) (where, *T_i_ = β_i_* log *n*). The total number of tests is given by *T*_1_ + *dT*_2_.

The defectives are expected to participate in a higher fraction of positive outcome tests than the normal items or the inhibitors. And, once the defectives are identified, tests of each one of them with rest of the items can be used to determine their associations. We show that this can be done non-adaptively as well. The decoding algorithm proceeds in two stages for both the non-adaptive as well as the adaptive pooling design. The first stage will identify the defectives from the outcome vectors **y** and **y**_1_ in the non-adaptive and adaptive pooling designs respectively, according to the fraction of positive outcome tests in which an item participates. The second stage will identify the inhibitors and their associations with the declared defectives using subsets of the outcome vector **y** in the non-adaptive pooling design and the outcome vectors **y***_̂u_1__*, **y***_̂u_2__* …, **y***_̂u_d__* in the adaptive pooling design.

Let us define the following notations^2^ with respect to the pools represented by **M***_NA_* and **M**_1_ which are eventually useful in characterizing the statistics of the different types of items that are used in the decoding algorithm.

*Notations:*

- ℐ(*u*) denotes the set of inhibitors that the defective *u* is associated with.
- ℱ*_uk_* denotes the event that none of the inhibitors associated with a defective *u_k_* appears in a test, given that the defective *u_k_* appears in the test.
- 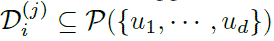 denotes the *j*^th^-set in the (arbitrarily) ordered set of all *i*-tuple subsets of the defective set denoted by 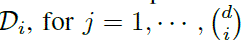, where *u_i_* denotes a defective and 𝒫 {(*u_1_, …, u_d_*)} denotes the power set of the set of defectives.
- 𝒟(*s*) denotes the defectives associated with the inhibitor *s* and its complement is given by 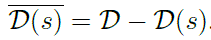.
- 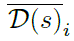 denotes the (arbitrarily) ordered set of all *i*-tuple subsets of the defective set 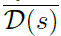 and the *j*^th^-set in 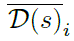 is denoted by 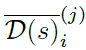.

*Example 2:* Realizations of the above notation for the association graph in Fig. 2 considered in Example 1 are given below. The inhibitor set is given by ℐ = {*s_1_* …, *s_r_*} and the defective set is given by 𝒟 = {*u_1_*, …, *u_d_*} with *r* = *d*.

- *ℐ(u)* for *u = u_1_* is given by ℐ(*u*_1_) = {*s*_1_}.
- ℱ_*uk*_ represents the event that the inhibitor *s*_1_ associated with the defective *u*_1_ does not appear in a test, given that the defective *u*_1_ appears in the test.
- Realizations of 𝒟_i_ for *i* =1, 2 are given by

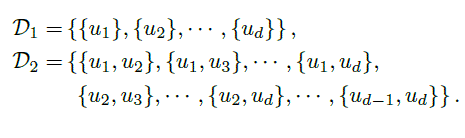

Realizations of 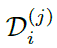 for (*i, j*) = (1,2) and (*i, j*) = (2,3) are given by

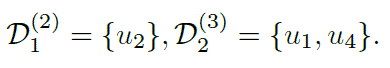
- 𝒟(*s*) for *s* = *s*_1_ is given by 𝒟(*s*_1_) = *u*_1_ and its complement is given by 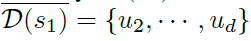.
- Realizations of 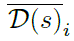 for *s = s_1_* and *i* = 1, 2

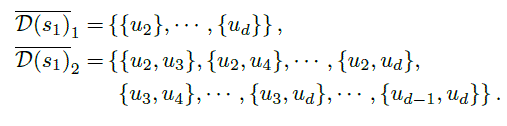

Realizations of 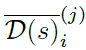 with *s = s_1_*, for (*i,j*) = (1, 2) and (*i, j*) = (2, 3) are given by

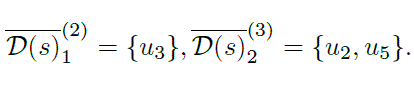

We now define the following statistics corresponding to the different types of items. The following statistics also hold good when **y**_1_ is replaced by **y**, as entries of both and **M***_NA_* and **M**_1_ have the same statistics.

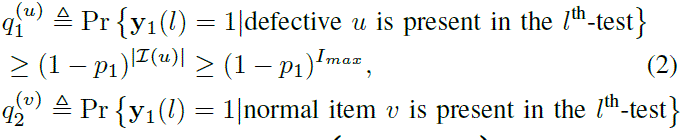

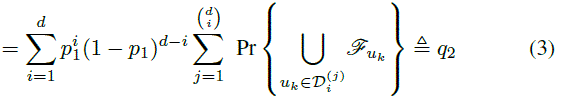

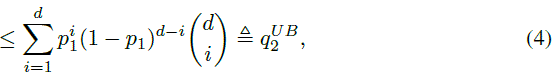

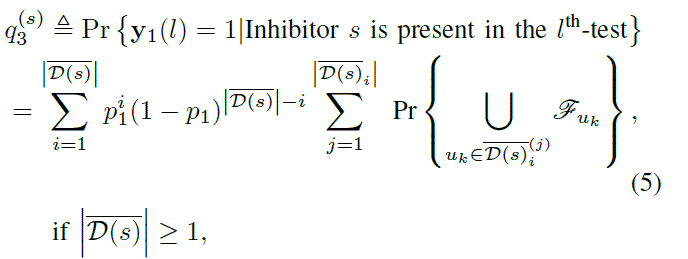

= 0, otherwise.

Since the outer and inner summations in (5) is over a subset of those in (3), max 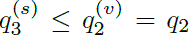. The upper bound in (4) follows from the upper bound of one on the probability terms of (3). In hindsight, the lower and the upper bounds can be easily obtained as follows. The lower bound on the positive outcome statistics for a defective item in (2) follows from the worst case statistics when all the inhibitors inhibit the expression of every defective. The upper bound on the statistics for a normal item in (4) follows from using the best case positive outcome statistics, in the absence of inhibitors, where the appearance of any defective gives a positive outcome. It is easy to see that positive outcome for an inhibitor in a test is less probable than that for a normal item. Denote the worst case negative outcome statistic for a defective by

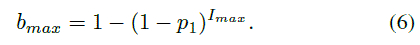

Denote the set of tests corresponding to outcome vector **y** in which an item *w_j_* participates by 𝒯*w_j_* (**y**) and the set of positive outcome tests in which the item *w_j_* participates by **𝒮***w_j_* (**y**), for *j* = 1, 2, …, *n*. The decoding algorithm is given as follows.

1. *Stage* 1 (*Identifying the defectives for <u>both non-adaptive and adaptive pooling designs):</u>* For the non-adaptive pooling design, if |**𝒮**_wj_ (**y**)| > 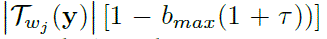 with *b_max_* as defined in (6), declare the item *w_j_* to be a defective. For the adaptive pooling design, we use the same criterion, replacing **y** by **y**_1_. Denote the number of items declared a.s defectives by *d̂* and the set of declared defectives by 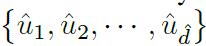. If *d̂* ≠ *d* declare an error. Denote the remaining unclassified items in the population by 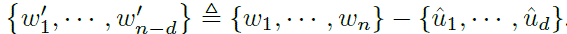
2. *Stage 2 (Identifying the inhibitors and their associations for <u>non-adaptive pooling design):</u>* Let 𝒫_k_ denote the sets of pools in **M**_*NA*_ that contain only the declared defective *û_k_* and none of the other declared defectives, for *k* = 1, …, *d*. Also, let the outcomes corresponding to these pools be positive. This means that the pools in *𝒫_k_* do not contain any inhibitor from the set ℐ(*û_k_*), which denotes the set of inhibitors associated with the item *û_k_* if *û_k_* is indeed a defective. Now, consider only the outcomes corresponding to these pools denoted by **y**_𝒫_1__ ⊂ **y**, …, **y**𝒫_*d*_ ⊂ **y**. The associations of the declared defectives are identified as follows.

- For each *k* = 1 to *d*, declare 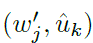 to be a nonassociated inhibitor-defective pair if 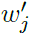 participates in at least one of the tests corresponding to the outcome vector **y** 𝒫_*k*_ and declare the rest of the items to be associated with *û_k_*. The items declared as non-associated for all *k* are declared to be be normal items. If 𝒫*_k_* = {ø} for some *k*, declare an error.
3. ***Stage*** 2 *(Identifying the inhibitors and their associations for <u>adaptive pooling design):</u>* Let 𝒮(**y***_û_k__*) denote the set of positive outcome tests corresponding to **y***_û_k__*, i.e., these pools do not contain any inhibitor from the set ℐ(*û_k_*) if *û_k_* is a defective.
  - For each *k =1* to *d*, declare 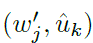 to be a nonassociated inhibitor-defective pair if 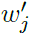 participates in at least one of the tests in the set 𝒮(**y***_ûk_*) and declare the rest of the items to be associated with *û_k_*.

The items declared as non-associated for all *k* are declared to be be normal items. If 𝒮(**y**_*û_k_*_) = {ø} for some *k*, declare an error.

*Remark 1: (Stage* 1) The first stage in the decoding algorithm, which is the same for both the non-adaptive and adaptive pooling design, is similar to the defective classification algorithm used in [13] for the 1-inhibitor model. The underlying common principle used is that there exists statistical difference between the defective items and the rest of the items. Hence, with sufficient number of tests, the defectives can be classified by “matching” the tests in which an item participates and the positive outcome tests. The items involved in a large fraction of positive outcome tests are declared to be defectives. A similar decoding algorithm was used in the classical group testing framework with noisy tests [22]. Here, the inhibitors of a defective item, if any, behave like a noise due to probabilistic presence in a test. The (worst case) expected number of positive outcome tests in which a defective participates is at least 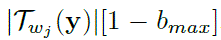. Like in [13], the Chernoff-Hoeffding concentration inequality [23] is used to bound the error probability and obtained the exact number of tests required to achieve a target (vanishing) error probability. *It is important to note that, a priori, it is not clear if a fixed threshold technique can sieve the defectives under worst case positive outcome statistics and the rest of the items under best case positive outcome statistics, with vanishing error probability*.

*Remark 2: (Stage 2)* In the IDG model, the inhibitors for each defective might be distinct. Hence, an inhibitor for one defective behaves as a normal item from the perspective of another defective. This defective-specific interaction is absent in the 1-inhibitor model. So, any inhibitor can be identified using any defective, i.e, an inhibitor's behaviour is defective-invariant in the 1-inhibitor model, which was exploited in identifying the inhibitors in [13]. Since each inhibitor's behaviour can be defective-specific in the IDG model, we need to identify the defectives first and then identify its associated inhibitors by observing the interaction of the other items with each of these defectives.

The following theorems state the values of the parameters *p_1_, p*_2_, and τ, and the scaling of the number of tests required for the proposed non-adaptive and adaptive pooling designs to determine the association graph with high probability. Similar results can be stated for the IDG-NSI model, by replacing ***I**_max_* by *r* in the following theorems.

*Theorem 1 (Non-adaptive pooling design):* Choose the pooling design matrix **M**_*NA*_ of size T*_NA_ × n* with its entries distributed as i.i.d. ℬ***(p_1_)*** with 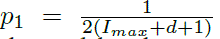 for the IDG-WSI model. Test the pools denoted by the rows of the matrix M_*NA*_ non-adaptively. The scaling of the number of tests sufficient to guarantee vanishing error probability (1) using the proposed decoding algorithm with 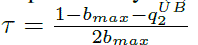 is given by 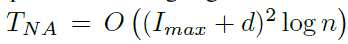, where 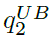 are defined in (4) and (6) respectively.

*Theorem 2 (Adaptive pooling design):* Choose the pooling design matrices **M**_1_ and **M**_2_ of sizes T*_1_ × n and T_2_ × n* with their entries distributed as i.i.d. ℬ*(p_1_)* and ℬ*(p_2_)* respectively, with 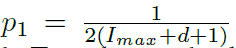 and 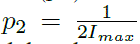 for the IDG-WSI model. Test the pools denoted by the rows of the matrices M_1_ non-adaptively and classify the defectives. Now, test each of the pools from M_2_ along with the *d* classified defectives individually. The scaling of the number of tests sufficient to guarantee vanishing error probability (1) using the proposed decoding algorithm with 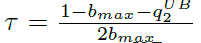 is given by *T_A_* = *T_1_ + dT_2_ = O (I_max_ d* log *n*), where 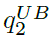 and *b_max_* are defined in (4) and (6) respectively.

The following sub-section constitutes the proof of the above theorems. The exact number of tests required to guarantee vanishing error probability for recovery of the association graph are also obtained. The proof is exactly the same for the IDG-NSI model, but replacing *I_max_* by *r*.

### A. Error Analysis of the Proposed Algorithm

As mentioned in Section II, we require that

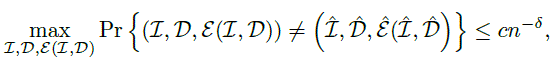

for some *c* > 0 and fixed *δ* > 0. We find the number of tests *T*_1_ required to upper bound the first stage error probability by *c*_1_*n*^−*δ*_1_^ and also the value of *T*_2_ required to upper bound the second stage error probability by *c*_2_*n*^−*δ*_2_^, for some constants *c*_1_ and *c*_2_. Finally, the values of *δ*_1_ and *δ*_2_ are chosen so that the total error probability is upper bounded by *cn*^−*δ*^, for some constant *c* and given *δ* > 0.

*1) Error Analysis of the First Stage:* Since the first stage of the decoding algorithm is the same for both the non-adaptive and adaptive pooling design, the bounds on the number of tests obtained below for adaptive pooling design applies for the non-adaptive pooling design also. The three possible error events in the first stage of the decoding algorithm for both non-adaptive and adaptive pooling design are given by

1. A defective is not declared as one.
2. A normal item is declared as a defective.
3. An inhibitor is declared as a defective.

Clearly, the defective that has the largest probability of a negative outcome, given by 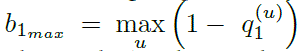, has the largest probability of not being declared as a defective. So, with *T*_1_ = *β*_1_ log *n*, the probability of the first error event for all the defectives can be upper bounded as

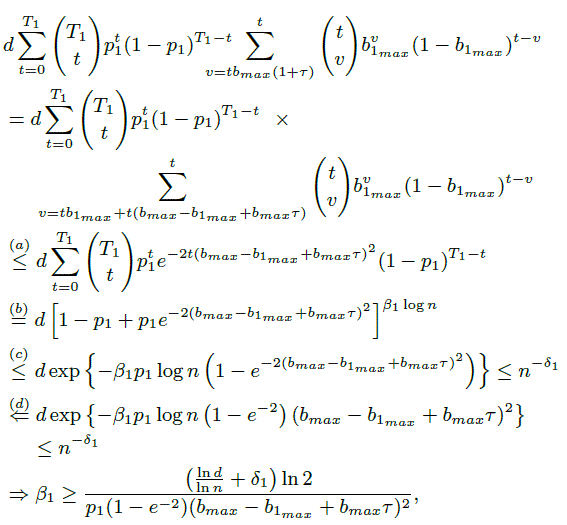

where (*a*) follows from Chernoff-Hoeffding bound [23]^3^, (*b*) follows from binomial expansion, (*c*) follows from the fact that 1 − *c* ≤ *e*^−*c*^, and (*d*) follows from the fact that (1 − *e*^−2*x*^2^^ ≥ (1 − *e^−2^*) *x*^2^, for 0 < *x* < 1. Using the fact that *b_1_max__ ≤ b_max_*, where *b_max_* is defined in (6), the following bound on *β*_1_ suffices.

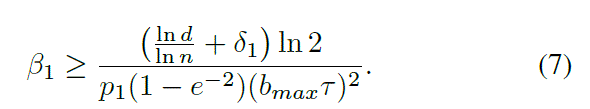

Similarly, to guarantee vanishing probability for the second error event (union-bounded over all normal items) and the third error event (union-bounded over all inhibitors), it suffices that

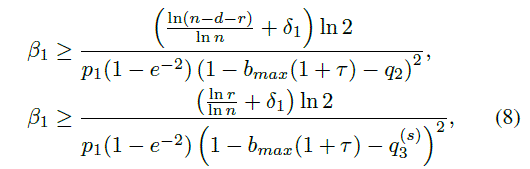

Since 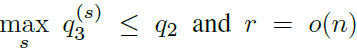, the bound in (8) is asymptotically redundant for all values of *τ*. So, substituting the upper bound on *q*_2_ defined in (4) by 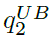, it suffices that

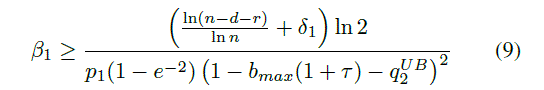

Now, the value of τ chosen to optimize the denominators of (7) and (9) is given by 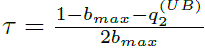. Therefore, we have

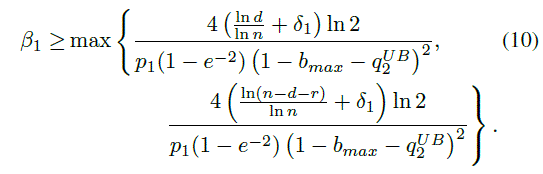

The term 1 − *b_max_* − *q*_2_ can be lower bounded as follows.

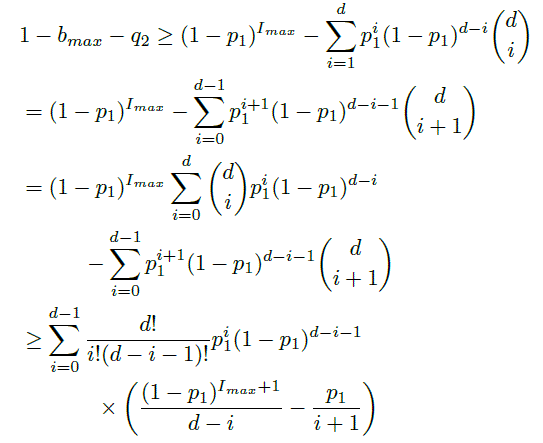

The above term can be non-trivially lower bounded if the last multiplicative term is positive for all *i*. So, it suffices if it is positive for *i* = 0. Using the lower bound (1 − *p*_1_)*^I_max_+^*^1^ ≥ 1 − (I_*max*_ + 1) *p*_1_, the above term at *i* = 0 is lower bounded as

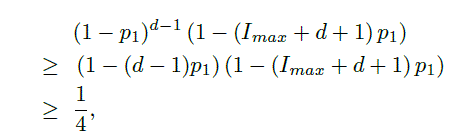

where the last lower bound follows by substituting *p*_1_ = 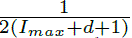. Hence, from (10) and using *r, d* = *o*(*n*), it suffices that

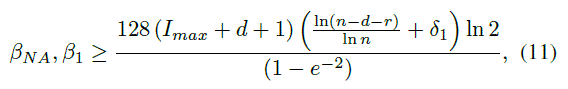

where *T_NA_* = *β_NA_* log *n*.

2) *Error Analysis of the Second Stage:* In the error analysis of the second stage, we assume that all the defectives have been correctly declared. Errors due to error propagation from the first stage shall be analyzed later.

*<u>Non-adaptive pooling design</u>:*

The only error event for the non-adaptive pooling design in the second stage is that there does not exist a set of pools 𝒫*_k_* such that they contain only the defective *u_k_* and none of its associated inhibitors ℐ(*u_k_*), and all its non-associated items appear in at least one of such pools. Denote this error event by 𝒰 (*u_k_*). Clearly, none of the inhibitors associated with *u_k_* will be declared as non-associated with *u_k_*. This follows from the definition of the set of pools 𝒫_*k*_ and the decoding algorithm.

The probability that a non-associated item appears along with a defective *u_k_*, but none of its associated inhibitors and none of the other defectives appear in a pool from **M**_*NA*_ is given by

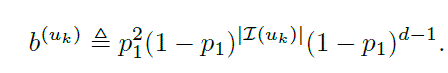

Now, probability of the error event is upper bounded by

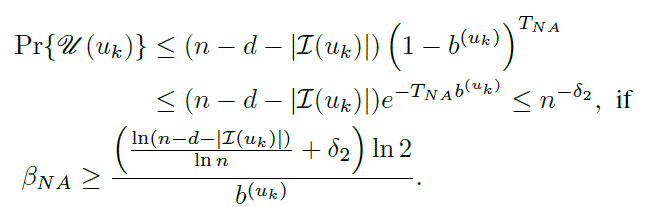

Since (1 ‒ *p*_1_)^|ℐ(*uk*)|^ ≥ (1 − *p*^*I*_*max*_^ ≥ (1 - *I_max_p*_1_) and (1 − *p*_1_)^*d*−1^ ≥ (1 − *dp*_1_), substituting for *p*_1_, it suffices that

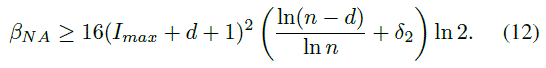

<u>*Adaptive pooling design:*</u>

Like in non-adaptive pooling design the only error event, denoted by ℰ*(u_k_*), is that items *w_j_* not associated with *u_k_* are declared as associated inhibitors, i.e., the item *w_k_* does not appear in any of the positive outcome tests *S(y_uk_*). Clearly, none of the inhibitors associated with *u_k_* will be declared as non-associated with *u_k_*.

Let *T*_2_ = *β*_2_ log *n*. The number of tests required to guarantee vanishing error probability for the error event ℰ*(u_k_)* is evaluated as follows. Let *w_j_* ∉ℐ(*u_k_)*. Define

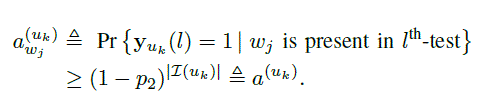

Now, we have

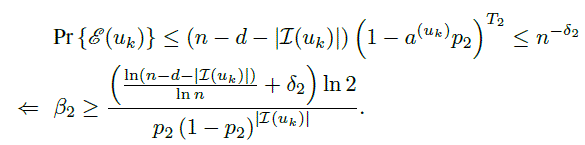

Using the fact that 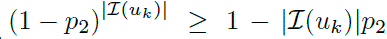, and substituting 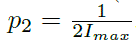, we have the following bound.

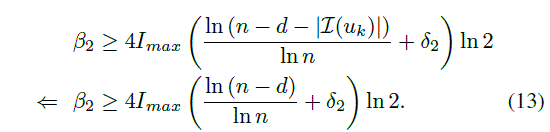

3) *Analysis of Total Error Probability*: Assuming that the target total error probability is *O(n*^−δ^), the values of δ_1_ and δ_2_ need to be determined. Towards that end, define the following events.

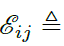 Event of declaring (*w_i_, w_j_), i = j*, to be an associated pair,

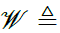 Event that at least one actual defective has not been declared as a defective.

Let ℰ denote the correct association pattern for some realization {ℐ, 𝒟}. Now, the total probability of error is given by

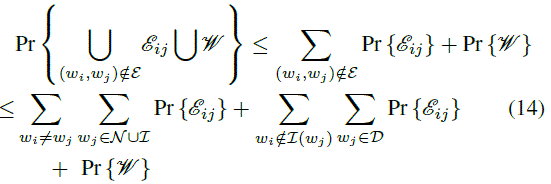

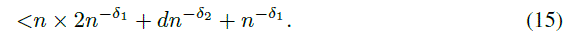

There are two possible ways in which the event ℰ*_ij_*, for (*w_i_,w_j_*) ∉ **ℰ**, can occur. One possibility is that the item *w_j_* has been erroneously declared as a defective in the first stage of the algorithm, and hence any item *w_i_* declared to be associated with *w_j_* is an erroneous association. The first term in (14) represents this possibility. The other possibility is that *w_j_* has been correctly identified as a defective, but the item *w_i_* is erroneously declared to be associated with *w_j_*. The second term in (14) represents this possibility. The last term accounts for the fact that a defective might be missed out in the first stage of the algorithm. Note that the other two terms do not capture this error event. Finally, (15) follows from the error analysis of the first and second stages of the decoding algorithm. Therefore, if the target error probability is *O*(*n*^−*δ*^), then choose *δ*_1_, *δ*_2_ = *δ* + 1.

Recall that the number of tests required for non-adaptive and adaptive pooling designs are given by *T_NA_ = β_NA_* log *n* and *T_A_* = *T*_1_ + *dT*_2_ = (*β*_1_ + *dβ*_2_) log *n* respectively. Therefore, from (11), (12), and (13) we have that *T_NA_ = O ((I_max_ + d)^2^* log *n*) and *T_A_* = *O (I_max_^d^* log *n*).

### B. Adaptation for the IDG-NSI Model

The only modification required in the pooling design and decoding algorithm proposed for the IDG-WSI model to adapt it to the IDG-NSI model is that *I_max_* is replaced by *r*. For the sake of clarity, we list the only changes below.

1) The pooling design parameters are chosen as *p*1 = 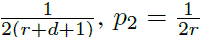.
2) In Stage-1 of the decoding algorithm the threshold for identifying the defectives is chosen as |𝒮w_j_ (**y**_1_)| > 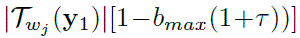, where *b_max_* = 1−(1−*p*_1_)^*r*^). Intuitively, this worst-case threshold corresponds to a scenario where every inhibitor inhibits every defective, i.e., the 1-inhibitor model.
3) The values of *β*_*NA*_, *β*_1_ and *β*_2_ are chosen as

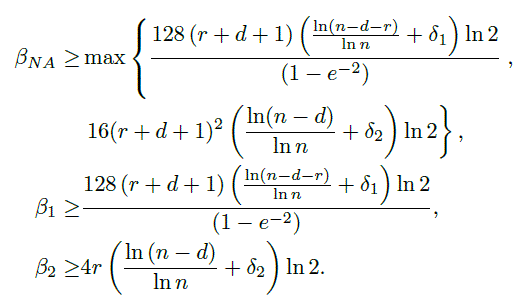

Hence, the total number of tests required for the IDG-NSI model scales as *T_NA_ = O* ((*r + d*)^2^ log *n*) for the nonadaptive pooling design and *T_A_* = *O*(*rd* log *n*) for the two-stage adaptive pooling design.

In the next section, lower bounds on the number of tests for non-adaptive and adaptive pooling designs are obtained.

## IV. Lower Bounds for Non-Adaptive and Adaptive Pooling Design

In this section, two lower bounds on the number of tests required for non-adaptive pooling designs for solving the IDG-NSI and IDG-WSI problems with vanishing error probability are obtained. one of the lower bounds is simply obtained by counting the entropy in the system and this lower bound also holds good for adaptive pooling designs. The other lower bound is obtained using a lower bound result for the 1-inhibitor model which is stated below.

### Theorem 3

(*Th*. 1, *[13]*): An asymptotic lower bound on the number of tests required for non-adaptive pooling designs in order to classify *r* inhibitors amidst *d* defectives and *n* − *d* normal items in the 1-inhibitor model is given by 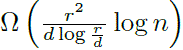, in the *d = o(r),r = o(n)* regime^4^.

### Theorem 4

An asymptotic lower bound on the number of tests required for non-adaptive pooling designs for solving the IDG-NSI problem with vanishing error probability for *r, d = o(n)* is given by

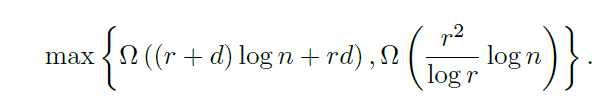

*Proof:* The proof for the first lower bound on the number of tests follows by lower bounding the number of possible combinations of the sets of defectives, inhibitors and associa tion patterns^5^.

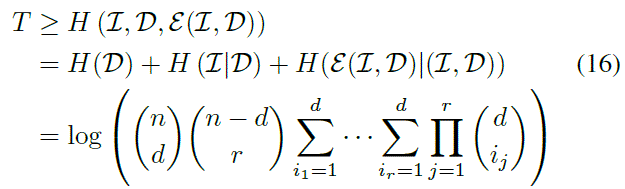

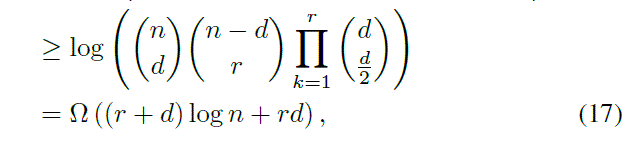

where *i_j_* denotes the number of defectives that the *j*^th^ inhibitor can be associated with, and the last step follows by using Stirling's lower bound 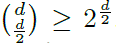. This lower bound is also valid for *adaptive pooling designs*.

The proof for the second lower bound for non-adaptive pooling designs is as follows. Assume that it's required to identify the inhibitors alone. Clearly, this requires lesser number of tests than the problem of identifying the association graph. Since the objective is to satisfy the error metric in (1), the error probability criterion

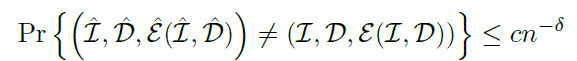

needs to be satisfied for all possible association patterns on all possible realizations of (ℐ, 𝒟). We choose one possible association pattern on some realization of (ℐ, 𝒟) given in Fig. 4 to evaluate the lower bound. Now, assume that a genie reveals the set of defectives 𝒟′ = {*u_2_, …, u_d_*} which are not associated with any of the inhibitors. A lower bound for this problem with side information from the genie is clearly a lower bound for the original problem. Note that the presence of any defective from the set 𝒟' in a pool always gives a positive outcome and hence, provides zero information for distinguishing the inhibitors from the rest of the items as the entropy of such outcomes is zero. Hence, the inhibitor identification problem for items with the association pattern as given in Fig. 4 is now reduced to the problem of identifying *r* inhibitors amidst *n − d* normal items and one defective item in the 1-inhibitor model, where *d* = *o*(*n*). For this problem, from Theorem 3, we have that the lower bound is given by 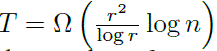 tests. Hence, this is also a lower bound on the number of tests required to identify the association graph with vanishing worst case error probability.

**Fig. 4.**
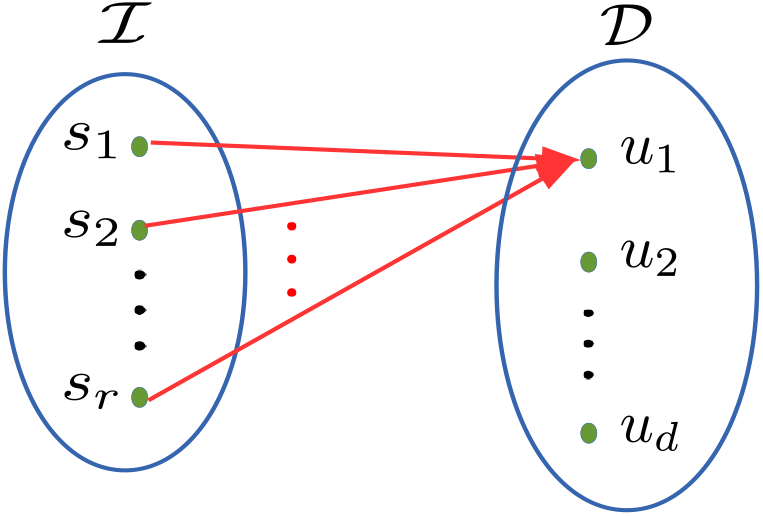
A possible association pattern for a possible realization of (ℐ, 𝒟) where a single defective is associated with all the inhibitors, but none of the other defectives are associated with any inhibitor.

The lower bounds for the IDG-WSI model are obtained in the following theorem. Since we are interested in asymptotic lower bounds, we assume that 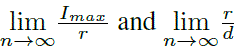 exist, and 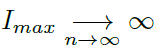

### Theorem 5

An asymptotic lower bound on the number of tests required for non-adaptive pooling designs for solving the IDG-WSI problem with vanishing error probability for *r, d = o(n)* is given by

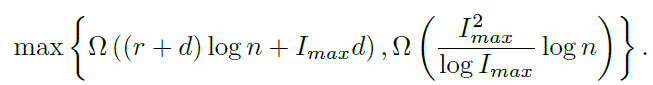

*Proof:* The first lower bound is obtained by lower bounding *H* (ℰ (ℐ, 𝒟)| (ℐ, 𝒟)) in (16) as follows. Two constraints need to be satisfied while counting the entropy of association pattern.

- *First constraint:* Minimum degree of a vertex in ℐ is one.
- *Second constraint:* Maximum degree of a vertex in 𝒟 is no more than *I_max_.*

We now consider the three possible cases below and show that in each of the cases the lower bound on the number of association patterns scales exponentially in *I_max_d.* Let *(c − 1)d < r ≤ cd*, for some positive integer *c*, and so *I_max_* ≥ *c*. Define 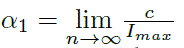 and 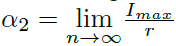.

Case 1: α_1_ < 1 and *α_2_ < 1*. There exist positive constants *β*_1_ < 1 and *β_2_ <* 1 so that *c* ≤ *β_1_I_max_* and *I_max_* ≤ *β_2_r,* ∀*n* ≥ *n*_0_. Define an association pattern, where each defective starting from *u_1_* is assigned a disjoint set of *c* inhibitors until every inhibitor is covered. Therefore we have, 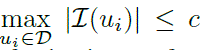. Since the first constraint is satisfied, each defective is now free to choose an association pattern so that 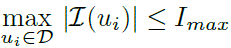. The number of such possible association patterns can be lower bounded by

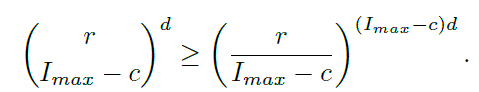

Thus, the entropy of association pattern in this case scales (asymptotically) as the logarithm of the above quantity, which is given by Ω*(I_max_^d^).*

*Case 2: α_1_ <* 1 and α_2_ = 1. There exist positive constants *β*_1_ < 1 and *β*_2_ ≤ 1 with *β*_2_ > *β*_1_ so that *c* ≤ *β_1_I_max_* and *I_max_ ≥ β*_2_ *r*, ∀*n* ≥ *n*_0_. So, we have *I_max_* − *c ≥ (*β*_2_ − *β*_1_)*r*, ∀*n* ≥ *n*_0_.* Using similar arguments as in Case 1, where after satisfying the first constraint, *β*_2_*r* − *c* inhibitors are chosen to associate with each defective, we now have that the entropy of association pattern in this case scales asymptotically as 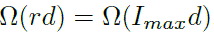

*Case* 3: *α_1_* = 1. Note that this case constitutes a large inhibitor regime with respect to the number of defectives (because *I_max_ → ∞*). There exists a positive constant *β*_1_ ≤ 1 so that *c* ≥ *β_1_I_max_, ∀n* ≥ *n*_0_. The number of ways of assigning each defective to a disjoint set of (*c* − 1) inhibitors is given by

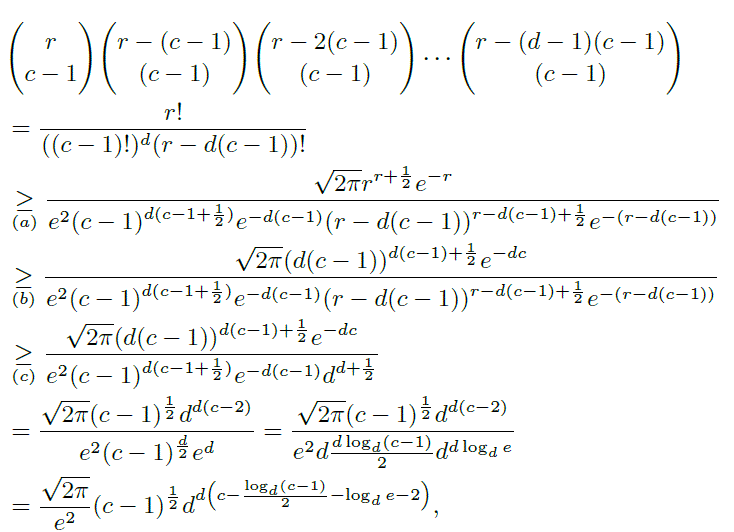

where (*a*) follows from Stirling's lower and upper bounds for factorial functions, (*b*) and (*c*) follow from the fact that *d*(*c* − 1) < *r* ≤ *cd.* Observe that the remaining *r* − *d*(*c* − 1) inhibitors can be assigned one each to one defective without violating the second constraint. Thus, the entropy of association pattern in this case scales asymptotically as Ω(*cd*) = Ω(*β*_1_*I_max_d*) = Ω (*I_max^d^_*).

The second lower bound is obtained as shown below. There can exist at least one defective *u*_1_ ∈ 𝒟 so that |ℐ(*u*_1_)| = *I_max_*. Consider an association pattern where ℐ(*u*_1_) ∩ ℐ(*u_k_*) = {ø}, for *u_k_* ∈ 𝒟, *k* ≠ 1, as depicted in Fig. 5. Now, we use a similar argument as in the proof of the second lower bound in Theorem 4. Let a genie reveal the inhibitor subset ℐ − ℐ(*u*_1_), the defective subset 𝒟 − *u*_1_ and their associations. Now, none of the items from the sets ℐ − ℐ(*u*_1_) and 𝒟 − *u*_1_ is useful in distinguishing the inhibitors in the set ℐ(*u*_1_ ) from the unknown defective and the normal items. This is because the entropy of an outcome is zero if the test contains some defective from 𝒟 − *u*_1_ but none of its associated inhibitors (which are only from the set ℐ − ℐ(*u*_1_)) as such a test outcome is always positive. The entropy of an outcome does not change if any of the inhibitors ℐ − ℐ*(u_1_)* with or without its associated defectives (which are only from the set 𝒟 − *u_1_*) is present in the test. Thus, the problem is now reduced to the 1-inhibitor problem of finding *I_max_* inhibitors amidst *n* − (*r* − *I_max_*) − (*d −* 1) normal items and one (unknown) defective. A lower bound on the number of non-adaptive tests for this problem is clearly a lower bound on the number of tests for the original problem of determining the association graph for the IDG-WSI model. Since *r, d* = *o*(*n*) and 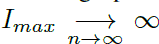, using Theorem 3, we get the lower bound 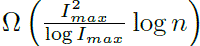.

**Fig. 5.**
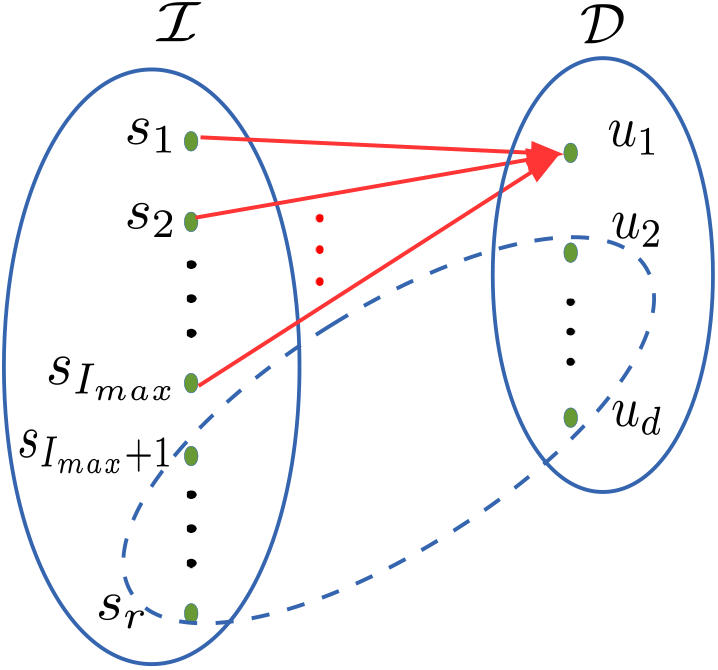
A possible association pattern where, without loss of generality, *u*_1_ is assumed to be a defective for which |ℝ(*u*_1_)| = *I_max_*. The set of inhibitors and defectives that are associated only among themselves are inside the dotted ellipse.

Thus, in the *d* = *O*(*I_max_*) and *d* = *O*(*r*) regimes, the upper bound on the number of tests for the proposed non-adaptive pooling design is away from the proposed lower bound for the IDG-WSI and IDG-NSI models by log *I_max_* and log *r* multiplicative factors respectively. But in the *I_max_* = *o*(*d*) and *r* = *o*(*d*) regimes, the upper bounds scale as *O*(*d*^2^ log *n*) which is far from the proposed lower bounds by atmost 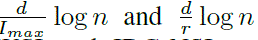 multiplicative factors for the IDG-WSI and IDG-NSI models respectively. For the proposed two-stage adaptive pooling design, the upper bound on the number of tests is away from the proposed lower bound by log n multiplicative factors for both the IDG-WSI and IDG-NSI models in all regimes of the number of defectives and inhibitors.

## V. Conclusion

A new generalization of the 1-inhibitor model, termed IDG model was introduced. In the proposed model, an inhibitor can inhibit a non-empty subset of the defective set of items. Probabilistic non-adaptive pooling design and a two-stage adaptive pooling design were proposed and lower bounds on the number of tests were identified. In the large inhibitor regime, the upper bound on the number of tests for the proposed non-adaptive pooling design is shown to be close to the lower bound, with a difference of a logarithmic multiplicative factor. However, in the small inhibitor regime, the upper bound is far away from the proposed lower bound. The nonadaptive pooling design made use of the principle of testing an isolated defective with the rest of the items to determine its associated inhibitors. This is the primary reason for the upper bound to be far away from the lower bound. It is not clear if this is a necessary condition for identifying all the associations. Tightening the lower bound, based on necessity of this condition, is an interesting problem.

For the proposed two-stage adaptive pooling design, the upper bound on the number of tests is close to the lower bound in all regimes of the number of inhibitors and defectives, the difference being logarithmic multiplicative factors. An interesting research direction is to include the *k*-inhibitor model, for unknown k, as a part of the association pattern in the IDG model, and derive lower and upper bounds on the number of tests for the resulting model. The proposed IDG model with noisy tests is also worth studying.

## Footnotes

1 The number of inhibitors, defectives and normal items are denoted by *r, d*, and *n – d* – *r* respectively.

2 From hereon, we reserve the notation *u* to represent a defective, *v* to represent a normal item and *s* to represent an inhibitor.

3 If the term (*b_max_ − b_max_ + b_max_τ) > 1*, then the probability of the error event under consideration is equal to zero.

4 Though Theorem 1 in [13] is stated for the classification of both the defectives and inhibitors in the 1-inhibitor model, it is also valid for classification of inhibitors alone.

5 The proof for the fact that the entropy of the system is a lower bound on the number of tests for both non-adaptive and adaptive pooling designs can be found in the appendix of [22].

